# Dietary quorum quenching AHL lactonase impairs the adaptation of commensal bacterium to host innate immunity

**DOI:** 10.1101/2025.02.05.636741

**Authors:** Rui Xia, Yuanyuan Yao, Delong Meng, Yajie Zhao, Shichang Xu, Ya Jin, Zhen Zhang, Yalin Yang, Chao Ran, Zhigang Zhou

## Abstract

Quorum sensing (QS) is the communication system of bacteria that depends on QS signals. Quorum quenching (QQ) enzymes degrade QS signals and are promising alternatives of antibiotics to treat bacterial infections. Here, we found that dietary QQ *N*-acyl homoserine (AHL) lactonase led to microbiota dysbiosis in zebrafish, with reduction of *Aeromonas* and enrichment of *Plesiomonas*. Through gnotobiotic zebrafish colonized with a minimal microbiota, we found that QQ-mediated microbial alteration relies on host Myd88 signaling and neutrophil elastase. Mechanistically, quorum quenching increased the susceptibility of commensal *Aeromonas* to host neutrophil elastase by impairing bacterial lateral flagellar system, leading to reduced colonization of *Aeromonas* and subsequent enrichment of *Plesiomonas* due to ecological competition of the two species. Together, we found that dietary QQ lactonase led to microbiota alteration by impairing the adaptation of commensal *Aeromonas* to host innate immunity, which provided novel insight in the role of quorum sensing in host-microbiota interaction.

## Introduction

Antibiotics play an important role in the treatment of infectious diseases in human and animals.^1^ However, in the past few decades, bacteria have become resistant against antibiotics through antibiotic resistance mutations^2^ and horizontal gene transfer among bacteria,^3^ leading to the emergence of multiple drug resistant (MDR) bacterial strains. Due to the antibiotics resistance issue, antibiotics alternatives are one of the most important topics in the research community.

Quorum sensing (QS) is a communication mechanism between bacteria that relies on secreted signaling molecules which regulate the coordinated behaviors of bacteria. Bacteria regulate gene expression by sensing changes in signaling molecules.^4^ There are various signaling molecules in bacteria, including auto-inducing signal type 1 (AI-1), type 2 (AI-2) and type 3 (AI-3). Among them, *N*-acyl homoserine lactone (AHL) (AI-1) is the most extensively studied quorum sensing signaling molecule in bacteria. QS regulates many biological processes, including virulence, competence, conjugation, resistance, motility and biofilm formation.^5^ Due to the wide range of phenotypes controlled by QS, QS inhibition, i.e., quorum quenching (QQ), has been one of the most promising alternatives to antibiotics.^6^ Quorum quenching can be achieved by different mechanisms: (i) inhibition of signal molecule synthesis; (ii) blockage of signal molecules-receptor interaction; (iii) degradation of signal molecules by QQ enzymes,^7^ which don’t enter cells or bind to a receptor, but hydrolyze signal molecules secreted by the bacteria.^8,9^ Unlike antibiotics, quorum quenching does not kill pathogens or restrict cell growth, but block bacterial communications and thus the virulence functions and behaviors.^9–11^ Therefore, QQ has been proposed as an eco-friendly and effective alternative to antibiotics for managing bacterial diseases and offers a potential solution to multidrug-resistant pathogens.^12,13^ Oral ingestion of QS inhibitors or QQ enzymes exhibited protective effect in different animal models against bacterial infection.^14–19^

Increasing evidence indicates that QS signal molecules mediated inter-species and inter-kingdom communication, which could influence the composition of the intestinal microbiota.^20,21^ For example, AI-2 favored Firmicutes and hindered Bacteroidetes, thus shaping the microbiota composition under antibiotics-induced dysbiosis in mice ^22^ Moreover, AI-2 could promote the colonization of *Lactobacillus rhamnosus* GG in neonatal mice.^23^ The effect of AHL on the composition of commensal microbes was also reported. *Hydra* modulated the colonization of its main commensal bacterium, *Curvibacter* sp, by modifying AHL molecules.^24^ The AHL signal molecule 3-oxo-C12:2 correlated with normobiosis in the human gut microbiota.^25^ Consistent with the role of QS molecules in regulating microbial composition, quorum quenching has been reported to affect the commensal microbiota. AHL lactonase that degrades the signal molecules altered the bacterial composition of oral cavity samples.^26^ Dietary supplementation of a QQ enzyme producing probiotic *Bacillus* sp. QSI-1 affected the composition of intestinal microbiota and decreased the abundance of commensal *Aeromonas* in fish.^27^ QS interference blocking the AI-2 communication affected the assembly process of oral biofilm microbiota, leading to delayed and fragile maturity of the microbiota.^28^ However, the mechanism of quorum quenching induced alteration in microbial composition, as well as the role of QS in the colonization of commensal bacteria, are largely unknown.

Zebrafish have been used as a powerful animal model to study the assembly of intestinal microbiota and host-microbiota interaction.^29^ Proteobacteria is abundant in zebrafish microbiota, and the commensal functions of Proteobacteria taxa such as *Aeromonas* have been reported in gnotobiotic zebrafish.^29^ In an attempt to use QQ enzyme to protect zebrafish from infection by pathogenic *Aeromonas veronii*, we unexpectedly observed that dietary QQ enzyme increased the infection of zebrafish by the pathogen, which was due to that the QQ enzyme induced microbial dysbiosis featured with an enrichment of commensal *Plesiomonas* and a reduction of commensal *Aeromonas*. We then utilized gnotobiotic zebrafish colonized with a minimal microbiota to investigate the underlying mechanism of the QQ enzyme induced microbiota alteration, and found that the QQ enzyme impaired the resistance of commensal *Aeromonas* to host-derived immune factors, leading to reduced colonization of *Aeromonas* and subsequent microbiota alteration.

## Results

### Quorum quenching enzyme AiiO-AIO6 inhibited biofilm formation and the hemolytic activity of pathogenic *Aeromonas veronii*

*A. veronii* is the most important pathogen for cyprinid fish septicemia.^30^ QS plays a vital role in biofilm formation and the production of virulence factors by virulent *A. veronii*, and AHL is a key signaling molecule that regulates QS system.^31^ The quorum quenching enzyme AiiO-AIO6 (hereafter referred to as AIO6) is an AHL lactone enzyme that degrades AHL by hydrolyzing the lactone ring. We assessed the ability of AIO6 to inhibit biofilm formation and aerolysin expression by the high-virulence strain *A. veronii* Hm091. As shown in Figure 1A, the addition of AIO6 reduced the quantity of biofilm at a dose dependent manner as determined by crystal violet staining (Figure 1A). Moreover, the inhibitory effect of AIO6 on biofilm formation was confirmed microscopically by testing bacterial biofilms on glass slides with AIO6 (Figure 1B). AIO6 did not affect bacterial growth in 24 h, indicating that AIO6 has no selective pressure to the bacterium (Figure 1C). Furthermore, AIO6 reduced the expression of virulence factor aerolysin (Figure 1D). Collectively, *in vitro* studies demonstrated that AIO6 inhibited the virulence and biofilm formation of the pathogenic *A. veronii*.

**Figure 1.**
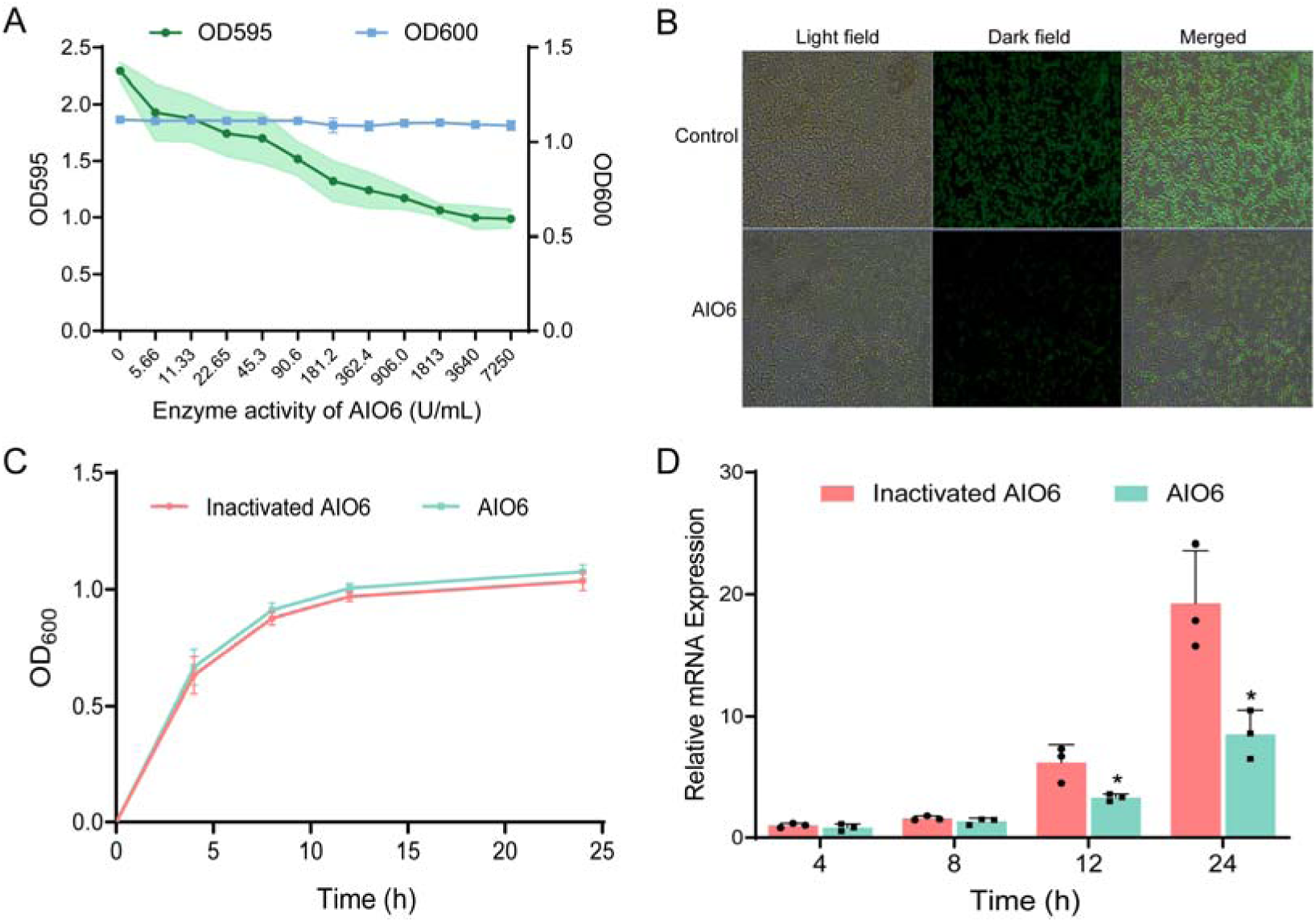
Inhibitory effects of quorum quenching enzyme AIO6 on biofilm formation and virulence gene expression of pathogenic *A. veronii* Hm091. (A) Growth curve of planktonic bacteria and biofilm production (n = 3). (B) Biofilm formation by microscopic validation. (C) Growth curve of *A. veronii* Hm091 treated with inactivated AIO6 and AIO6 (n = 3). (D) Transcription levels of aerolysin were determined by *q*PCR assays. Data are expressed as the mean ± SD. \**p* < 0.05, \*\**p* < 0.01, \*\*\**p* < 0.001.

### Dietary AIO6 led to gut microbiota dysbiosis and increased the susceptibility of zebrafish to pathogenic *A. veronii*

Immersion challenges experiment was performed to evaluate the protection effect of AIO6 against *A. veronii* Hm091 infection in zebrafish. Unexpectedly, the survival of zebrafish fed with AIO6 was decreased compared with control (Figure 2A). To explore the underlying mechanism, we first tested the direct effect of AIO6 on GF zebrafish. GF zebrafish were immersed with different concentrations of AIO6, and were challenged by the pathogen. As shown in Figure 2B, there were no significant changes in the mortality of GF zebrafish among control and AIO6 treated groups, ruling out the possibility that AIO6 directly impairs the antibacterial capacity of zebrafish. To evaluate the involvement of intestinal microbiota, 16S rRNA gene sequencing was conducted. The overall structure of intestinal microbiota in AIO6 fed zebrafish significantly differed from that of control group, as analyzed by principal coordinate analysis (PCoA) (Figure 2C). Moreover, taxon-based analysis revealed marked changes in the composition of intestinal microbiota. At the phylum level, the abundance of Fusobacteria was significantly decreased in AIO6 group, and Proteobacteria was markedly increased (Figure 2D and 2F). At the genus level, AIO6 group featured a significant decrease in the abundance of *Aeromonas* and an enrichment of *Plesiomonas* (Figure 2E and 2F). The abundance of *Cetobacterium* was also decreased in AIO6 group. To investigate the role of gut microbiota in the AIO6-mediated impairment of disease resistance in zebrafish, we fed zebrafish with Control, AIO6 or AIO6 supplemented with antibiotics. The effect of AIO6 on the disease resistance was abrogated by antibiotic treatment (Figure 2G), suggesting the involvement of intestinal microbiota. To confirm the causal relationship between AIO6-associated microbiota and disease resistance, we transferred the intestinal microbiota of Control and AIO6 groups to GF zebrafish, and tested the disease resistance (Figure 2H). The results showed that the phenotype of disease resistance in GF zebrafish colonized with microbiota from different groups were consistent with the donor zebrafish, suggesting that the dietary AIO6 impaired disease resistance of zebrafish through the intestinal microbiota.

**Figure 2.**
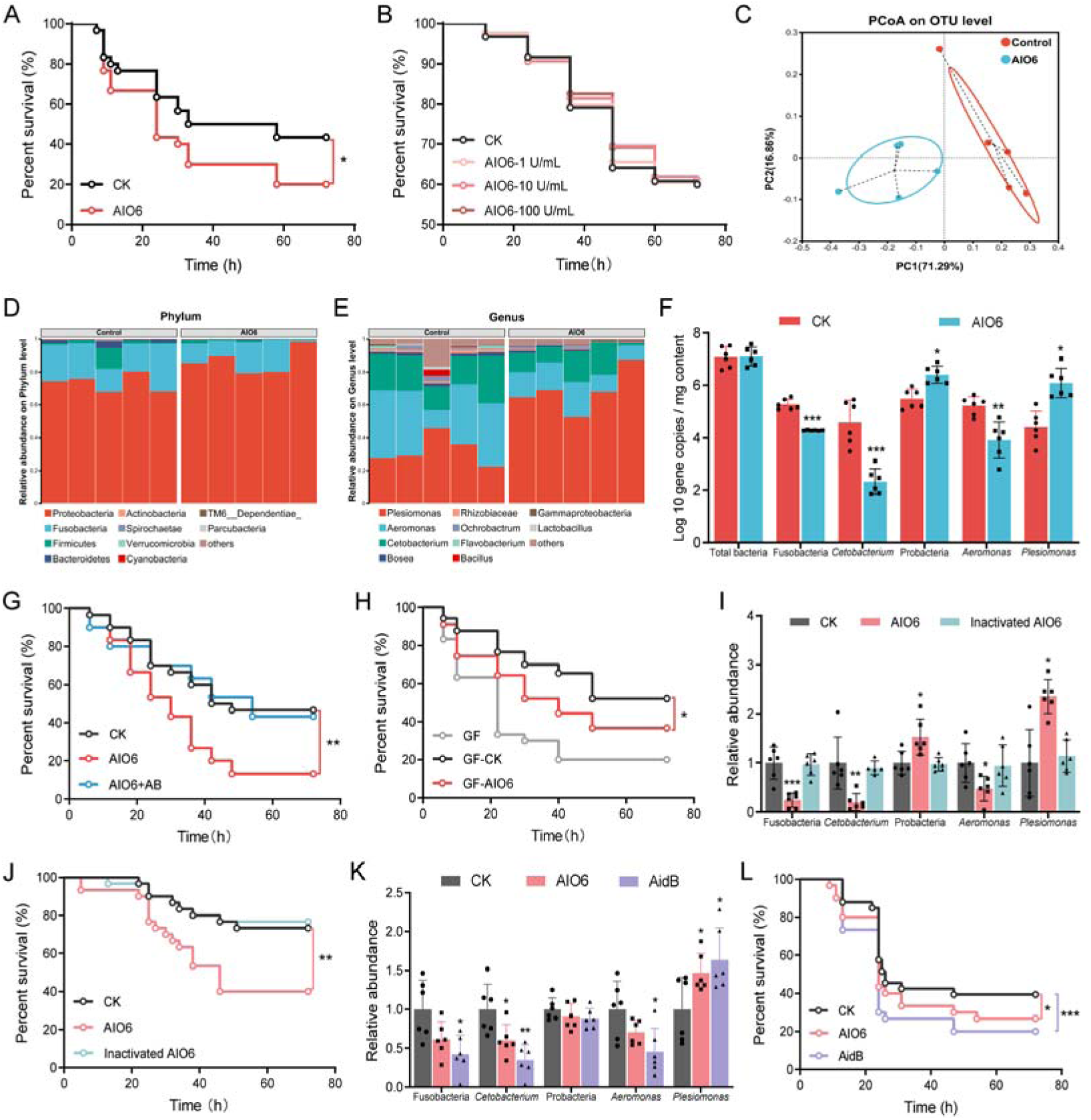
Dietary AIO6-induced microbiota dysbiosis increased pathogen susceptibility of zebrafish. Adult zebrafish (2-month-old) were fed with control, AIO6, inactivated AIO6 or AidB diet for 1 week. (A) Survival curve of adult zebrafish (n = 30). (B) Survival curve of GF zebrafish infected with *A. veronii* Hm091 (n = 90). (C) Principal coordinate analysis (PCoA) of the intestinal microbiotaby weighted UniFrac distance. (D and E) The relative bacterial abundance of gut microbiota of adult zebrafish fed control or AIO6 diet at the phylum (D) and genus (E) levels. (F) Bacterial gene copy numbers (n = 6, pool of 3 fish per sample). (G) Survival curve of adult zebrafish fed control, AIO6 and AIO6 + AB diets after *A. veronii* Hm091 challenge (n = 30). (H) Survival curve of GF zebrafish colonized with gut microbiota from adult zebrafish fed control and AIO6 diets (n = 90). (I) Relative abundance of bacterial taxa analyzed by *q*PCR (n = 6, pool of 3 fish per sample). (J) Survival curve of adult zebrafish fed control, AIO6 or heat-inactivated AIO6 diet after *A. veronii* Hm091 challenge (n = 30). (K) Relative abundance of bacterial taxa analyzed by *q*PCR (n = 6, pool of 3 fish per sample). (L) Survival curve of adult zebrafish fed control, AIO6 or AidB diet after *A. veronii* Hm091 challenge (n = 30). Data are expressed as the mean ± SD. \**p* < 0.05, \*\**p* < 0.01, \*\*\**p* < 0.001.

To confirm that the effect of AIO6 was mediated through its QQ enzymatic activity, we feed zebrafish with AIO6 and heat-inactivated AIO6, and found that heat-inactivation blocked the effect of AIO6 on the intestinal microbiota and disease resistance of zebrafish (Figure 2I and 2J). To investigate if the effect of AIO6 can be extended to other AHL lactone enzymes, AidB, an AHL lactonase belonging to the metallo-β-lactamase superfamily, was used.^32^ Results showed that dietary AidB also significantly reduced the abundance of *Aeromonas* and *Cetobacterium*, while increased the abundance of *Plesiomonas* (Figure 2K). Moreover, AidB increased pathogen susceptibility similarly as AIO6 (Figure 2L). These results indicate that AIO6 affects microbiota composition and disease resistance of zebrafish via the QQ enzymatic activity, and this phenotype is extensive in AHL lactonases.

### A minimal microbiota including *Aeromonas* and *Plesiomonas* recapitulated AIO6-induced alteration of the microbiota in gnotobiotic zebrafish

We attempted to verify the effects of AIO6 on the microbiota composition and disease resistance in zebrafish larvae. The results showed that the effect of AIO6 on the microbiota composition and disease resistance in zebrafish larvae was consistent with that in adult zebrafish (Figure 3A and 3B). In order to establish a simplified model to investigate the mechanism of AIO6-mediated microbiota alteration, we chose commensal *Aeromonas veronii*, *Plesiomonas shigelloides* and *Cetobacterium somerae* (hereafter referred to as *Aeromonas*, *Plesiomonas*, and *Cetobacterium*, respectively) as representatives of the microbiota, because of their prominence and considerable abundance change in response to AIO6. Gnotobiotic zebrafish were colonized with the three bacterial species and were fed AIO6. The results showed that the abundance of *Aeromonas* and *Cetobacterium* decreased, while the abundance of *Plesiomonas* increased after feeding AIO6, which were consistent with the microbial alteration observed in complex microbiota (Figure 3C). To further simplify the model, gnotobiotic zebrafish were colonized with a random pair of the three species. The results showed that AIO6 significantly reduced the abundance of *Aeromonas* while increased the abundance of *Plesiomonas* in zebrafish colonized with the pair of *Aeromonas* and *Plesiomonas* (Figure 3D-3F), while AIO6-induced bacterial alteration in zebrafish colonized with the other two pairs (*Cetobacterium* + *Aeromonas*, *Plesiomonas* + *Cetobacterium*) did not accord with the alteration observed in the three species model, indicating that *Aeromonas* and *Plesiomonas* dictated the AIO6-mediated microbial alteration, while the change in the abundance of *Cetobacterium* was probably the subsequent result. We also evaluated the effect of AIO6 feeding on the abundance of bacterium in mono-colonized gnotobiotic zebrafish, and observed no effect of AIO6 on the level of *Aeromonas* (Figure 3G) or *Plesiomonas* (Figure 3H). Therefore, a minimal microbiota consisting of *Aeromonas* and *Plesiomonas* was used to evaluate the mechanism of AIO6-induced microbial change. Analysis of *in vitro* interactions between *A. veronii* and *P. shigelloides* showed nutritional competition between the two species (Figure 3I and 3J), suggesting that AIO6-mediated alteration in the abundance of the two species involved microbe-microbe ecological competition.

**Figure 3.**
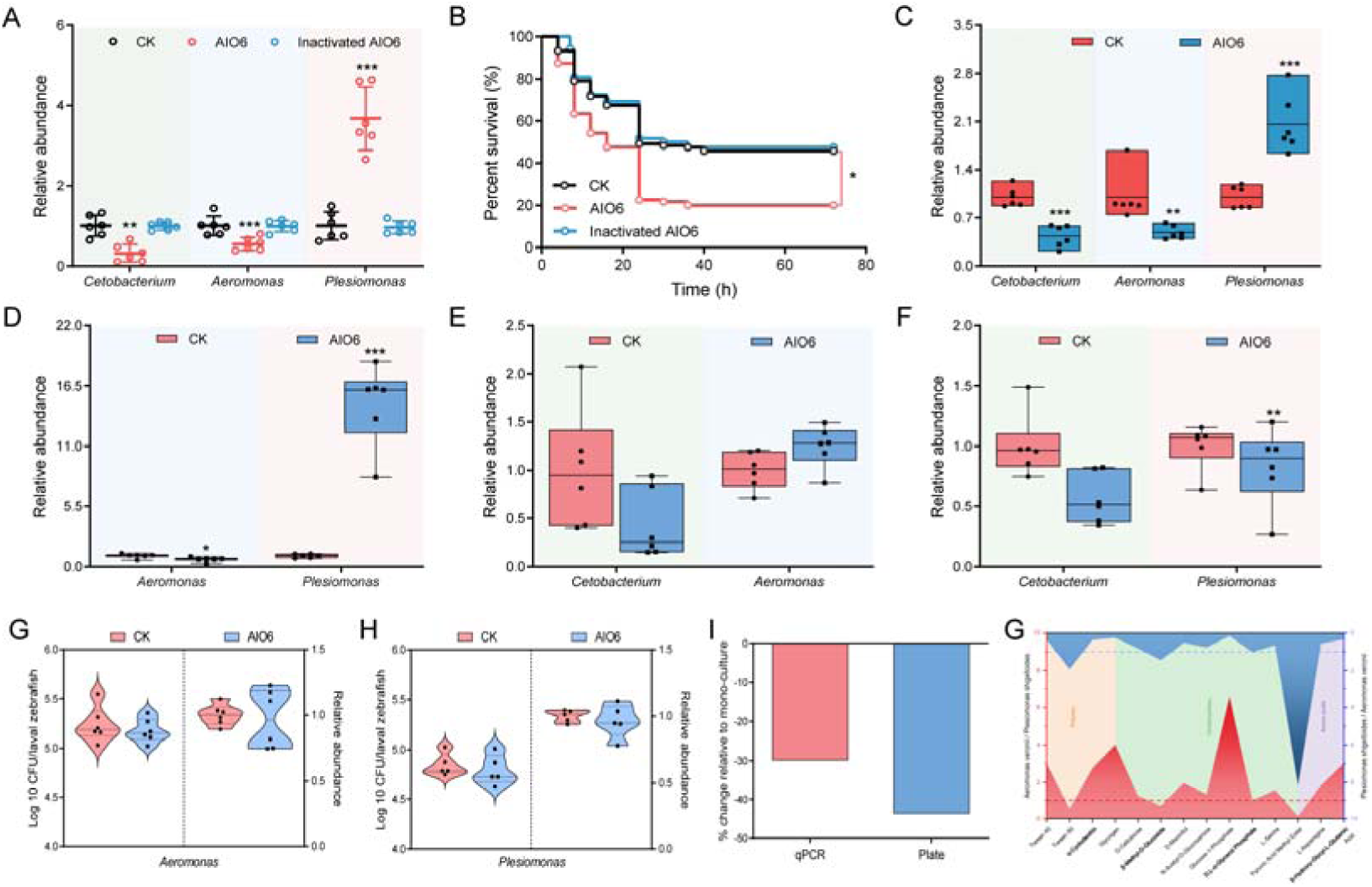
A minimal microbiota including *Aeromonas* and *Plesiomonas* recapitulated AIO6-induced alteration of the microbiota in gnotobiotic zebrafish. Zebrafish larvae were fed with control, AIO6 or inactivated AIO6 diet for 1 week. (A) Relative abundance of bacterial taxa in the gut of larval zebrafish (n = 6, pool of 20 fish per sample). (B) Survival curve of zebrafish larvae (n = 90). (C) The relative abundance of *Aeromonas*, *Plesiomonas*, and *Cetobacterium* in gnotobiotic zebrafish after AIO6 feeding. (D-F) The relative abundance of bacterial species in gnotobiotic zebrafish colonized with a random pair of the three species (n = 5 or 6, pool of 20 zebrafish per sample). (G and H) The relative and CFU abundance of bacterial species in mono-associated gnotobiotic zebrafish (n = 6, pool of 10 zebrafish per sample). (I) Bar plots showing the percentage change in the *in vitro* growth of co-culture of commensal *Aeromonas* and *Plesiomonas* in comparison to the monoculture. (J) Heatmap showing substrate utilization profile of the commensal *Aeromonas* and *Plesiomonas* using Ecoplate (BIOLOG). Data are expressed as the mean ± SD. \**p* < 0.05, \*\**p* < 0.01, \*\*\**p* < 0.001.

### AIO6-mediated alteration in commensal bacteria relies on the functioning of host-derived neutrophil elastase

No significant difference in the microbiota composition was observed by AIO6 treatment under *in vitro* condition (Figure S1), suggesting that the mechanism of AIO6-mediated effect involved host factors. To investigate if AIO6-induced microbiota alteration involved host immunity, gnotobiotic zebrafish colonized with *Aeromonas* and *Plesiomonas* were treated with prednisolone, a steroid immunosuppressant^33^ to suppress the immunity. The results showed that prednisolone treatment blocked the effect of AIO6 on the abundance of the two species, supporting the involvement of host immunity in AIO6-mediated effect on the microbiota (Figure 4B). Furthermore, we evaluated the effect of AIO6 on intestinal microbiota in zebrafish treated with vivo-morpholino directed to Myd88. Myd88 is an adaptor for innate immune receptors, including the Toll-like receptors. Therefore, Myd88 morphant zebrafish are immunodeficient. Interestingly, we observed no difference in the abundance of *Aeromonas* and *Plesiomonas* between AIO6-fed group and control in Myd88 morphant zebrafish (Figure 4C), further corroborating that the AIO6-mediated action on the commensal bacteria involved host immunity.

**Figure 4.**
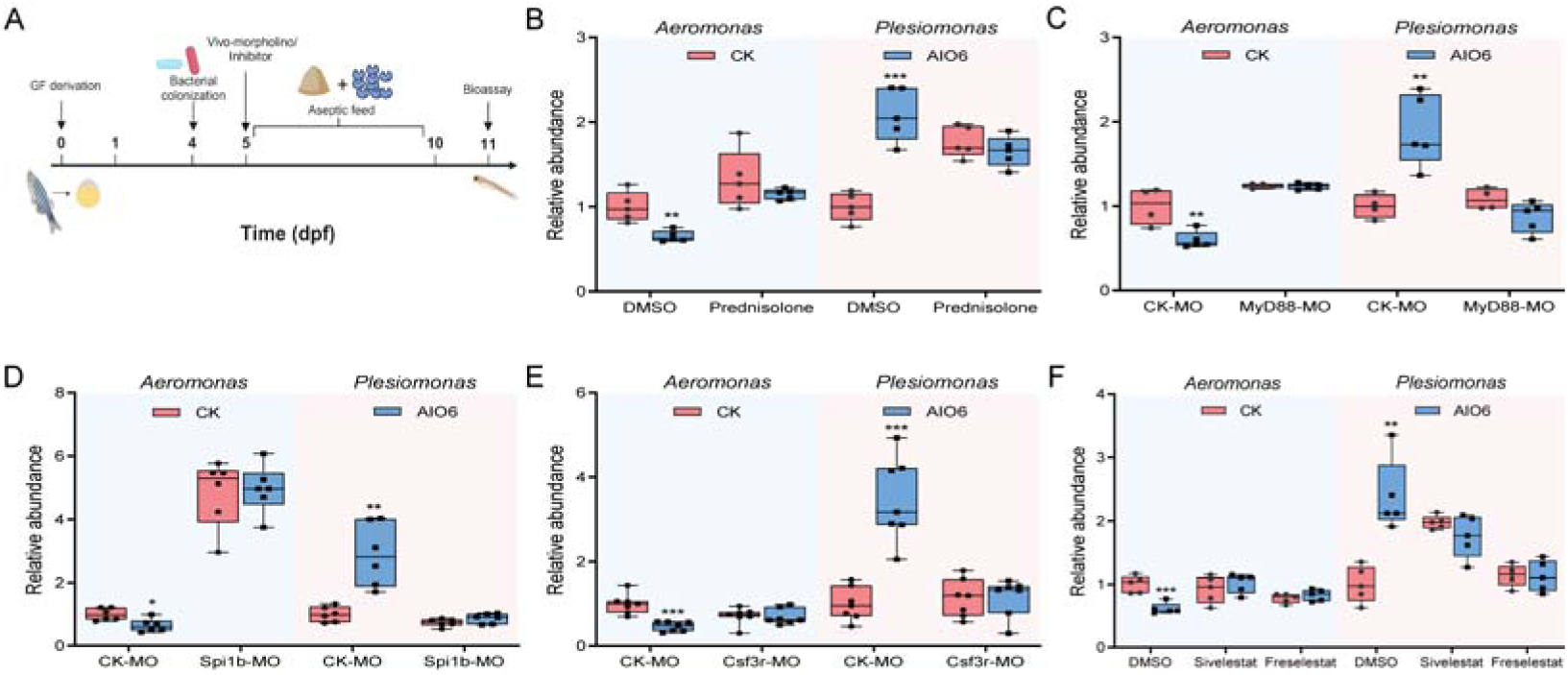
AIO6-mediated alteration in commensal bacteria relies on the functioning of host innate immunity. (A) Schematic representation for gnotobiotic zebrafish experiment (created using Biorender.com). (B-F) Effect of prednisolone (B), MyD88 MO (C), or Spi1b MO (D), or Csf3r MO (E), or NE inhibitors (F) on the relative abundance of bacterial species upon AIO6 feeding in gnotobiotic zebrafish colonized with *Aeromonas* and *Plesiomonas* (n = 4 or 6, pool of 20 zebrafish per sample). Data are expressed as the mean ± SD. \**p* < 0.05, \*\**p* < 0.01, \*\*\**p* < 0.001.

Then we investigated the cellular immunity involved in AIO6-mediated effect on the microbiota. At the larval stage of zebrafish, cellular immunity is composed only of myeloid cells, and neutrophils and macrophages are the main effector cells. We firstly depleted both neutrophils and macrophages using Spi1b (pu.1) MO. Spi1b MO blocked the effect of AIO6 on the abundance of *Aeromonas* and *Plesiomonas* (Figure 4D), indicating that the effect of AIO6 on intestinal bacteria depends on the function of myeloid cells. Knockdown of Csf3r affect the neutrophil population of zebrafish larvae.^34^ Thus, we attempted to deplete neutrophils using Csf3r MO to identify the effector cells. We found that Csf3r MO blocked the regulatory effect of AIO6 on the commensal bacteria (Figure 4E). In summary, these results indicated that AIO6-mediated effect on the abundance of intestinal bacteria involved the function of neutrophils.

As the first line of defense in the body, neutrophils are the first cells to reach the site of inflammation and release a variety of proteases involved in the inflammatory response.^35^ Neutrophil elastase (NE), a serine proteinase, which mainly secreted by neutrophils, plays an important role in immune defense.^36,37^ We treated zebrafish with Sivelestat and Freselestat, highly specific NE inhibitors,^38^ to evaluate the potential involvement of neutrophil elastase. The result showed that both Sivelestat and Freselestat blocked AIO6-induced alteration in the abundance of *Aeromonas* and *Plesiomonas* (Figure 4F). In addition, we inhibited innate immunity in conventional zebrafish larvae or gnotobiotic zebrafish colonized with a triple consortium including *Aeromonas*, *Plesiomonas*, and *Cetobacterium*, and found the effect of AIO6 on the commensal bacteria also depended on host Myd88 signaling and neutrophil elastase (Figure S2), corroborating that the results obtained from the bi-species minimal microbiota can be extended to more complex system.

We searched the *Plesiomonas shigelloides* genome sequence data (The genomic data have been deposited in NCBI under Bioproject PRJNA1205539) and found the *Plesiomonas* strain does not harbor an AHL QS system, excluding that *Plesiomonas* can directly respond to AIO6. To evaluate the specific role of *Plesiomonas* in AIO6-mediated alteration of the consortium, we treated *Aeromonas*-colonized GF zebrafish with cell lysate or cell free supernatant (CFS) of *Plesiomonas* and evaluated the abundance of *Aeromonas* in response to AIO6 feeding. The results showed that AIO6 significantly reduced the abundance of *Aeromonas* in mono-associated gnotobiotic zebrafish treated with *Plesiomonas* CFS (Figure 5A), indicating that AIO6-mediated reduction of *Aeromonas* requires some secreted substance(s) of *Plesiomonas*, but not the alive bacterium. Bacterial extracellular vesicles (EV) have been reported as important mediator of host-microbe interaction.^39,40^ To investigate the involvement of *Plesiomonas*-derived EVs in the AIO6-mediated effect on *Aeromonas*, we treated *Plesiomonas* with GW4869, a neutral sphingomyelinase inhibitor that impairs the release of EVs,^41^ to inhibit EVs secretion. We then observed that GW4869 abrogated the effect of *Plesiomonas* CFS in mono-associated zebrafish (Figure 5B), indicating that the *Plesiomonas*-derived EVs contributed to AIO6-induced reduction of *Aeromonas* in gnotobiotic zebrafish. Furthermore, we found that heat and proteinase K treatment blocked the effect of *Plesiomonas* CFS on AIO6-mediated reduction of *Aeromonas* in mono-associated gnotobiotic zebrafish (Figure 5C), indicating that the secreted effector mediated the action was probably a protein. Furthermore, we found that Myd88 and Csf3r knockdown, as well as Sivelestat, blocked the effect of AIO6 on the abundance of *Aeromonas* in mono-associated gnotobiotic zebrafish treated with *Plesiomonas* CFS (Figure 5D-5F), which was consistent with the results in gnotobiotic zebrafish colonized with bacterial consortium. In addition, *Plesiomonas* CFS had no effect on the growth of *Aeromonas* in the presence and absence of AIO6 *in vitro* (Figure 5G), excluding a direct effect on *Aeromonas* and suggesting an effect on host immunity. Together, these results support that *Plesiomonas* derived EVs were required for the functioning of innate immunity that was involved in the AIO6-mediated alteration of commensal bacteria.

**Figure 5.**
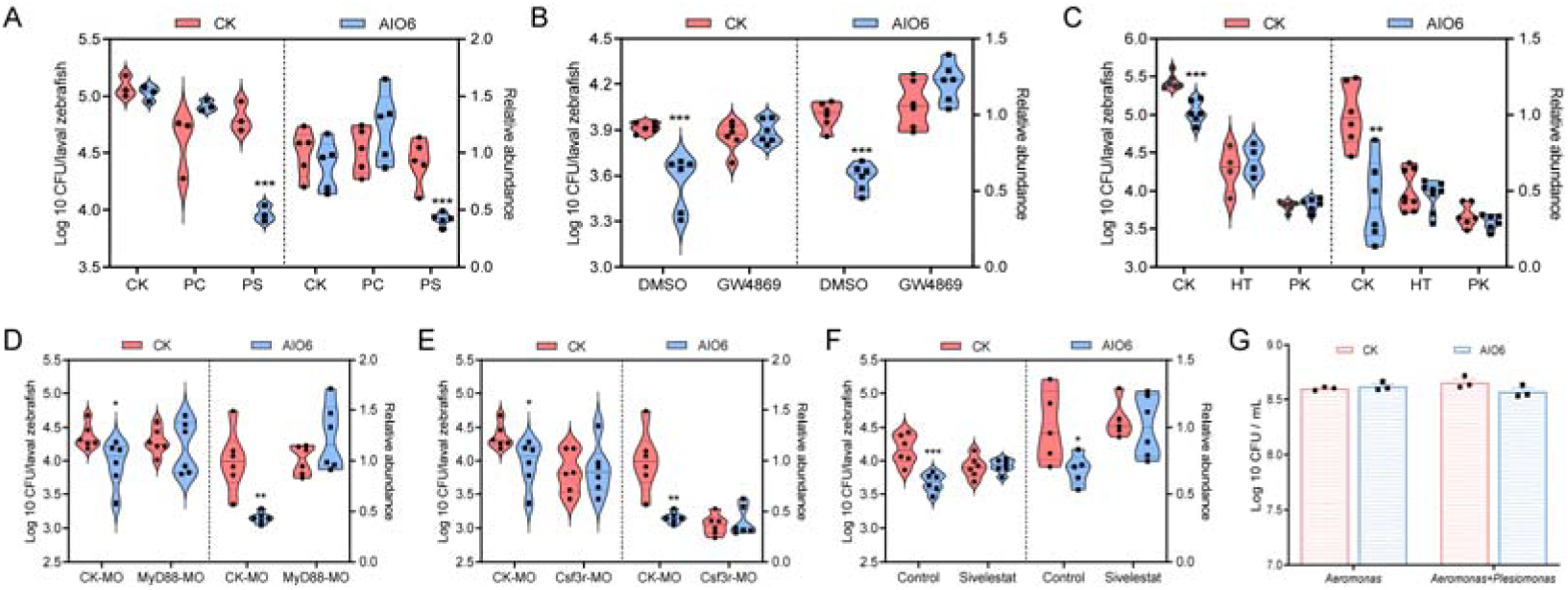
*Plesiomonas* contributes to AIO6-induced reduction in *Aeromonas* colonization through its extracellular vesicles (EVs) (A) The abundance of *Aeromonas* in GF zebrafish mono-colonized with *Aeromonas* and treated with the cell lysate (PC) or cell free supernatant of *Plesiomonas* (PS) (n =3 or 5, pool of 20 zebrafish per sample). (B) The abundance of *Aeromonas* in GF zebrafish mono-colonized with *Aeromonas* and treated with the cell free supernatant of *Plesiomonas* or GW4869-treated *Plesiomonas* (n =6, pool of 10 zebrafish per sample). (C) Effect of heat- or proteinase K-treated cell free supernatant of *Plesiomonas* on the abundance of *Aeromonas* in GF zebrafish mono-colonized with *Aeromonas* (n = 4 or 6, pool of 20 zebrafish per sample). HT, the cell free supernatant of *Plesiomonas* was heated at 100L for 5 min. PK, the cell free supernatant of *Plesiomonas* was treated with proteinase K at a concentration of 25 μg/ml at 70L for 10 min. (D-F) Effect of MyD88 MO (D), or Csf3r MO (E), or Sivelestat (F) on the relative and CFU abundance of *Aeromonas* in mono-colonized gnotobiotic zebrafish treated with cell free supernatant of *Plesiomonas* (n = 5 or 6, pool of 10 zebrafish per sample). (G) The growth of *Aeromonas* in the presence and absence of cell free supernatant of *Plesiomonas* under *in vitro* condition. Data are expressed as the mean ± SD. \**p* < 0.05, \*\**p* < 0.01, \*\*\**p* < 0.001.

### AIO6 increased the susceptibility of commensal *Aeromonas* to host-derived neutrophil elastase by impairing the lateral flagellar system

Several phenotypes in *Aeromonas* have been shown to be controlled by their quorum sensing systems, such as biofilm formation, flagella, and secretion system.^42–45^ We then investigated the mechanism underlying the effect of AIO6-mediated reduction of *Aeromonas* in mono-associated gnotobiotic zebrafish treated with *Plesiomonas* CFS. Firstly, we knocked out the quorum sensing system of the commensal *Aeromonas* strain, including AHL synthase gene *ahyI* and receptor gene *ahyR*, and investigated the AIO6-mediated effect on abundance of *Aeromonas* mutants. The results showed that AIO6 did not affect the colonization of Δ*ahyI* and Δ*ahyR* (Figure 6A). Consistently, AIO6 did not affect the colonization of Δ*ahyI* and Δ*ahyR* (Figure S3) in gnotobiotic zebrafish colonized with *Aeromonas* and *Plesiomonas*. These results indicated that AIO6 inhibited some downstream functions mediated by QS system to reduce bacterial colonization. To further determine the downstream functions that affect the colonization of *Aeromonas* in gnotobiotic zebrafish following AIO6 feeding, we knocked out genes related to the secretion system (*exeMN*, *ascV*, and *vasH*), flagellar system (*rpoN* and *lafK*), or bacterial adhesion and biofilm formation (*minD*). The results showed that the effect of AIO6 was abrogated for *Aeromonas* mutants deficient in the genes related to the flagellar system (Figure 6C), while the abundance of other *Aeromonas* mutants was still reduced in response to AIO6 feeding (Figure 6B) in gnotobiotic zebrafish, suggesting that AIO6 reduced the colonization of *Aeromonas* by influencing the flagellar system.

**Figure 6.**
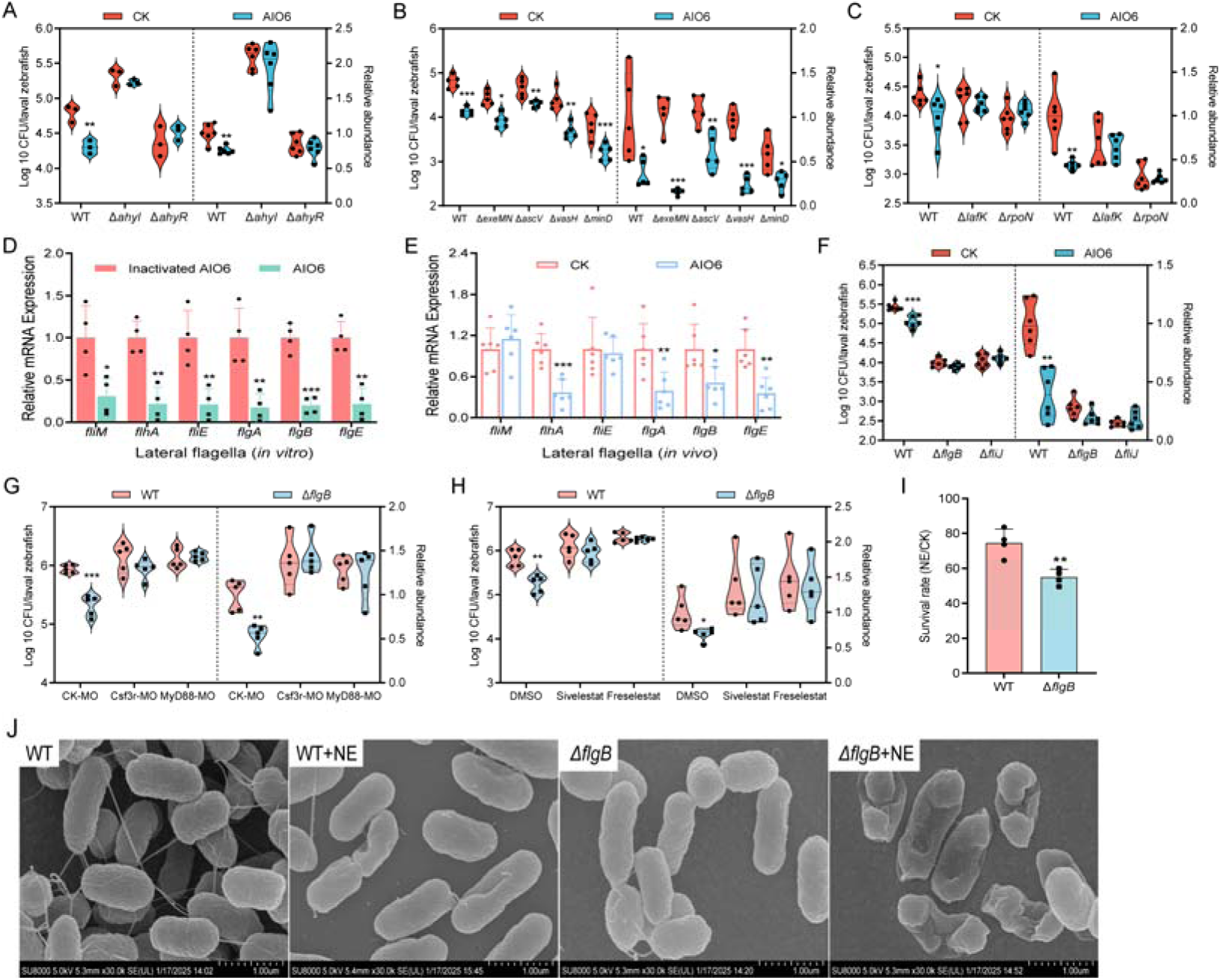
AIO6 impaired the adaptation of *Aeromonas* to host neutrophil elastase by inhibiting the lateral flagellar system. (A-C) The relative and CFU abundance of *Aeromonas* WT strain and mutant strains deficient in the genes related to AHL synthesis and reception (A), secretion system or biofilm formation (B), and flagellar system (C) in mono-associated gnotobiotic zebrafish in response to AIO6 feeding (n =6, pool of 20 zebrafish per sample). (D and E) Gene expression of the lateral flagellar system of commensal *Aeromonas* strain co-cultured with AIO6 under *in vitro* condition (D, n = 4) or in mono-associated gnotobiotic zebrafish fed AIO6 (E, n = 6, pool of 25 zebrafish per sample). (F) The relative and CFU abundance of *Aeromonas* WT strain and mutant strains deficient in the genes related to lateral flagellar system in mono-associated gnotobiotic zebrafish in response to AIO6 feeding (n =6, pool of 20 zebrafish per sample). (G and H) The relative and CFU abundance of WT strain versus Δ*flgB* mutant in mono-associated gnotobiotic zebrafish. (I) Viability of WT strain or Δ*flgB* mutant after incubation with neutrophil elastase. (J) The morphological differences between WT strain and Δ*flgB* mutant after incubation with neutrophil elastase as measured by SEM (scale bar, 1.0 μm). Data are expressed as the mean ± SD. \**p* < 0.05, \*\**p* < 0.01, \*\*\**p* < 0.001.

We then evaluated the effect of AIO6 on the flagellar system of commensal *Aeromonas*. AIO6 exhibited significant inhibitory effects on the expression of genes related to the lateral flagellar and polar flagellar systems under *in vitro* condition (Figure 6D and S4A). We also evaluated the effect of AIO6 on *Aeromonas* in mono-associated gnotobiotic zebrafish. The results showed that AIO6 mainly inhibited gene expression in the lateral flagellar system, and had no effect on the polar flagella (Figure 6E and S4B). Then we knocked out genes related to the lateral flagellar system, including *flgB* (Rod) and *fliJ* (Export/assembly), and tested the response of the mutants to AIO6 in gnotobiotic zebrafish. As shown in Figure 6F, AIO6 had no effect on the colonization of *Aeromonas* mutants deficient in the genes related to lateral flagellar system, indicating that dietary AIO6 reduced that abundance of *Aeromonas* by interfering with the lateral flagellar system.

In line with that AI06-mediated impairment of lateral flagellar system led to reduced *Aeromonas* colonization, *Aeromonas* Δ*flgB* mutant showed lower colonization versus the wild-type (WT) strain in mono-associated gnotobiotic zebrafish treated with *Plesiomonas* CFS. Interestingly, we found that Csf3r and Myd88 knockdown, as well as NE inhibitor (Sivelestat and Freselestat), abrogated the differential colonization of WT stain and Δ*flgB* mutant in gnotobiotic zebrafish (Figure 6G and 6H), supporting that the lateral flagellar system plays a role in the adaptation of commensal *Aeromonas* to host innate immunity (with neutrophil elastase as the immune effector). To directly investigate whether the lateral flagellar system contributes to NE susceptibility of commensal *Aeromonas*, we incubated WT strain and Δ*flgB* mutant with human NE, and bacterial viability was monitored by plating serial dilutions. Compared with WT strain, Δ*flgB* mutant showed significantly lower cell viability after NE treatment (Figure 6I). Furthermore, we cultured WT strain and Δ*flgB* mutant strain with or without NE, and the reactions were processed for scanning electron microscopy (SEM). WT strain showed concave pits and slightly deformed morphology after NE treatment. However, the surface structure of Δ*flgB* mutant was significantly damaged, and the bacterial cells showed signs of cracking (Figure 6J). Together, the results suggest that AIO6 increased the susceptibility of commensal *Aeromonas* to host-derived neutrophil elastase by impairing the lateral flagellar system, leading to reduced colonization of commensal *Aeromonas* and a subsequent enrichment of *Plesiomonas* due to ecological competition of the two species (Figure 3I and 3J).

## Discussion

Previous studies on bacterial QS were mostly conducted in the context of pathogen infections.^46,47^ The intestinal microbiota is a multi-species consortium with high bacterial cell density. As a way of bacterial communication, QS is believed to play a role in the microbe-microbe and host-microbe interactions in the microbiota.^48,49^ For instance, commensal *Bacillus* has been reported to eliminate *Staphylococcus aureus* by interfering with quorum sensing of the pathogen.^15^ Quorum sensing of commensal *Serratia ureilytica* drives outer membrane vesicle biogenesis and promotes its gut colonization in mosquito.^50^ Nevertheless, the studies on the role of QS signals in host-microbe interactions in the context of microbiota are only in the beginning, and our knowledge in this subject is limited. In this study, by dietary supplementation of a quorum quenching lactonase, we observed a role of QS in the colonization of commensal *Aeromonas* in zebrafish, and the concomitant role in the assembly and composition of the microbiota. In particular, QS signaling of commensal *Aeromonas* maintains lateral flagellar system of the species, which promotes adaptation to host immunity and gut colonization. Our results are consistent with previous study,^50^ providing another evidence that supports the important role of QS in gut colonization of commensal bacteria.

The inter-bacterial interactions hold important clues in shaping microbiota diversity and mediating responses of microbiota to exogenous factors, such as diets and pathogens.^51^ In previous studies, commensal bacterial consortia have been proved to decolonize bacterial pathogens through nutrient competition.^52,53^ In addition to ecological interaction among bacteria, the commensal microbiota can mediate colonization resistance through host immunity. Commensal Bacteroidetes protect against *Klebsiella pneumoniae* colonization through IL-36 and macrophages.^54^ Commensal bacteria, specifically clostridial Firmicutes and Bacteroidetes, inhibited *Candida albicans* colonization by activation of intestinal HIF-1α and LL-37.^55^ In this study, dietary AIO6 impaired the resistance of *Aeromonas* to host-derived neutrophil elastase, leading to reduced colonization of *Aeromonas*. Nutrient competition between *Aeromonas* and *Plesiomonas* exists due to the overlap in nutrient-utilization profiles, leading to an enrichment of *Plesiomonas* subsequent to the reduction of *Aeromonas*. Our results revealed the complexity of microbiota assembly and shift, which involved both host-microbe and microbe-microbe interactions.

Studies have shown an active interaction between host immune system and commensal microbiota. Commensal bacteria can regulate different aspects of innate and adaptive immunity, including IgA production, mucus secretion, and production of innate immune effectors such as antimicrobial peptides (AMPs).^56^ Reciprocally, host immunity exerts control and regulation on the microbiota. For example, the antimicrobial lectin RegIIIγ prevents commensal bacterial colonization of the intestinal epithelial surface and thus maintains the spatial segregation of microbiota and host.^57^ The antimicrobial peptide α-defensins have been shown to regulate the composition of the intestinal microbiota.^58^ The control of host immunity on specific microbial species has also been reported. For instance, T lymphocytes control intestinal microbial composition by suppressing the growth of *Vibrio* species in zebrafish,^59^ with unknown effector molecule(s). In this study, we found that neutrophil elastase can control the growth of commensal *Aeromonas*. Neutrophil elastase has been reported to kill bacterial pathogens after infection.^60^ The results in our study suggest the role of neutrophil elastase in control of intestinal microbiota, which sheds light on novel aspect of the function of this innate immune factor. Robinson et al. reported that evolved commensal *Aeromonas* showed competitive advantage in colonization relative to the ancestor strain that are specific in WT but not in Myd88^-/-^ zebrafish, suggesting a selective force on commensal *Aeromonas* colonization conferred by Myd88-mediated host immunity.^61^ Our results suggest neutrophil elastase as a potential effector molecule of the Myd88-mediated host immunity in the control of *Aeromonas* colonization, as Myd88 knockdown also blocked AIO6-mediated reduction of *Aeromonas* in gnotobiotic zebrafish (Figure 5D). The potential link and pathway between Myd88 and intestinal neutrophil elastase deserve further investigation. Quorum quenching has been proposed as promising alternatives of antibiotics to treat bacterial infections. Although the results in this study were obtained in an oral ingestion model, injection of small molecule quorum quenching inhibitors might have similar influences on the intestinal microbiota. Together, our results suggest that quorum quenching might influence the colonization of commensal bacteria sharing QS signal with the target pathogen, leading to a shift in the microbiota and thus an influence on health and disease.

## Resource availability

### Lead contact

Further information and requests for resources and reagents should be directed to the lead contact, Zhigang Zhou.

### Materials availability

Materials in this work are available from the lead contact.

### Data availability

The source data supporting the conclusions of this article are provided with this paper. Raw sequences have been deposited in NCBI as Bioproject # PRJNA1209212 and PRJNA1205539. This paper does not report original code. Any additional information required to reanalyze the data reported in this paper is available from the lead contact upon request.

## Acknowledgements

This work was supported by the National Natural Science Foundation of China (No.31925038, No.32122088, No.32303027) and the Postdoctoral Fellowship Program of China Postdoctoral Science Foundation (Grant Number GZC20241950).

## Author contributions

*A.* C. R. and Z. Z. designed the experiments; R. X. conducted the experiment and analyzed the data; Y. Y., D. M., Y. Z., S. X., Y. J., Z. Z. and Y. Y. helped with the experiments or provided the reagents; R. X. and C. R. prepared the manuscript.

## STAR★Methods

### Key resources table

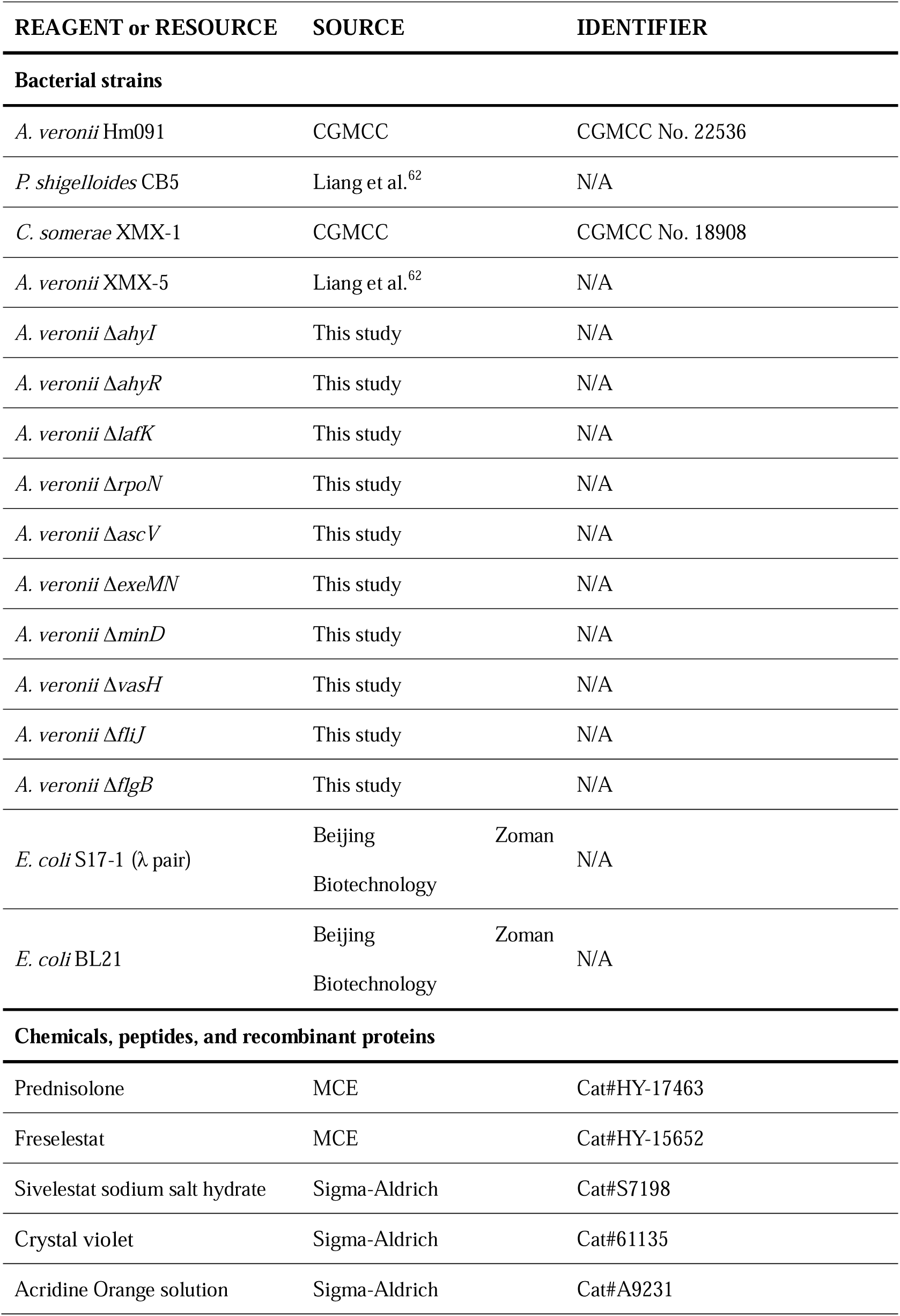
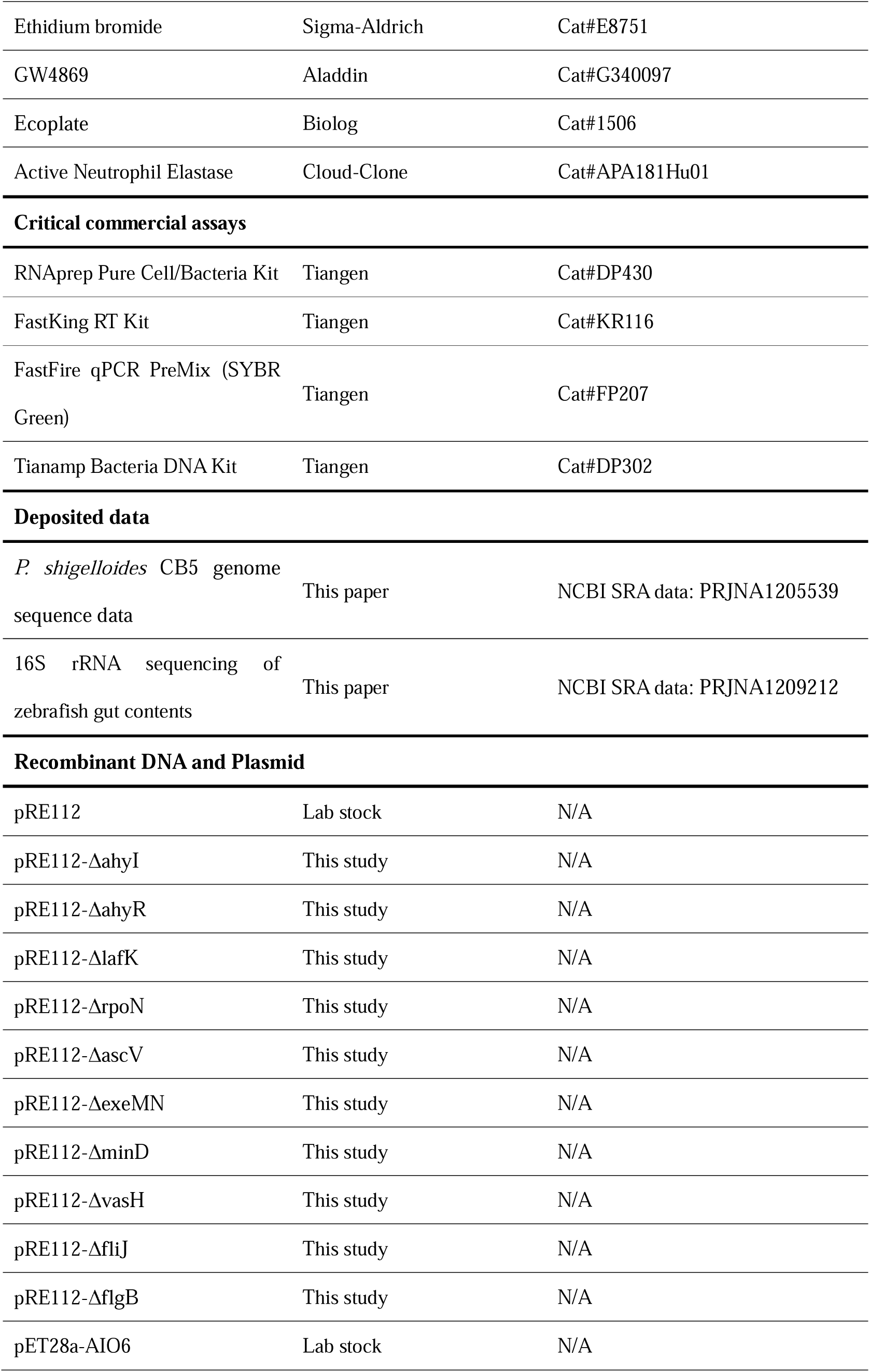

### Experimental model details

#### Zebrafish husbandry and experimental diets

All experiments in this study were approved by the Feed Research Institute of the Chinese Academy of Agricultural Sciences Animal Care Committee under the auspices of the China Council for Animal Care (Assurance No. 2018-AF-FRI-CAAS-003).

Adult zebrafish (Tu line) were bred in the fish lab of Institute of Feed Research of Chinese Academy of Agricultural Sciences, Beijing, China. Zebrafish were reared in recirculating aquaculture tanks under controlled conditions (25-28°C, a 14:10 light/dark cycle) and fed two times per day.

For the feeding trial, 2-month-old zebrafish (n = 4 tanks / group, 18 fish per tank) were fed with the basal diet and experimental diets supplemented with AIO6 (10 U/g diet), inactivated AIO6 (by boiling at 100L for 10 min), or AidB (10 U/g diet) for 1 week (Table S1). For antibiotics feeding, zebrafish were fed with diet supplemented with antibiotic mixture (Polymyxin B 2.5 g/kg diet and Neomycin 3.3 g/kg diet) for 1 week (Table S2). Zebrafish larvae were reared in gnotobiotic zebrafish medium (GZM) at 28°C to 4 days post fertilization (dpf). Larvae were allocated randomly to tanks with 30 mL of GZM and at 30 larvae per tank at 4Ldpf. The larvae started to feed at 5Ldpf, and were fed with the experimental diets once a day for 6 days (Table S3).

At the end of experiments, zebrafish were challenged with pathogenic bacteria *Aeromonas veronii* Hm091. Hm091 was cultivated overnight, then inoculated into LB medium and cultured at 37□ for 18 h. Adult zebrafish were immersed in culture water at a concentration of 2.0 × 10^7^ CFU/mL, and zebrafish larvae were immersed in GZM at a concentration of 1 × 10^7^ CFU/mL. Zebrafish mortality was observed for 3 days.

#### Germ-free and gnotobiotic zebrafish experiments

Germ-free (GF) zebrafish were prepared following established protocols as described previously.^63^ The diets for GF or gnotobiotic zebrafish were sterilized by irradiation with 20 kGy gamma ray. GF zebrafish were allocated randomly to tanks with 30 mL of GZM at 30 larvae per tank. Transfer of gut microbiota from adult zebrafish to GF zebrafish was performed according to the method described by Rawls et al.^64^ In many cases, GF zebrafish were associated at 4 dpf with single bacterial species or a simplified microbiota consisting of two or three species. *C. somerae* XMX-1 was incubated anaerobically at 28°C for 16 h, while *A. veronii* XMX-5 and *P. shigelloides* CB5 were cultured at 30°C for 16 h. Then, the bacterial cells were collected by centrifugation at 4, 000 g for 5 min. The collected bacterial cells were washed three times and resuspended in phosphate buffered saline (PBS, *p*H 6.8). GF zebrafish were colonized at 4 dpf with bacterium at a final concentration of 10^6^ CFU/mL. Gnotobiotic zebrafish were fed control or AIO6 diet for 6 days before bioassay. When applicable, zebrafish were treated with DMSO, 25 μg/mL prednisolone, 20 μM Sivelestat, 15 μM Freselestat at 5 dpf, or were treated with vivo morpholino targeting genes related to innate immunity at 1 or 5 dpf.

#### Bacterial strains and culture conditions

The pathogenic *A. veronii* Hm091 was isolated from the liver of diseased carp. Commensal strains *Cetobacterium somerae* XMX-1, *A. veronii* XMX-5 and *Plesiomonas shigelloides* CB5 were isolated from the intestine of adult zebrafish (*Danio rerio*). *A. veronii*, *P. shigelloides*, and *Escherichia coli* were cultured in LB liquid medium or plated on LB agar. *C. somerae* was cultured in Gifu Anaerobic Medium (GAM) liquid medium or plated on GAM agar. The vector pRE112 was used as the suicide vector for gene knockout. AIO6 (QQ AHL lactonase from *Ochrobactrum* sp. M231) was expressed in *E. coli* BL21 cells from pET28a(+) plasmid, and purified from cell lysates using Ni-NTA columns. When necessary, appropriate concentrations of antibiotics were prepared: 100 μg/mL ampicillin (Amp), 50 μg/mL Kanamycin, and 30 μg/mL chloramphenicol (Cm) were used for the positive selection.

### Method details

#### Biofilm Formation Measurement

The inhibition of biofilm formed by *A. veronii* Hm091 was tested using microtiter plate assay^65^ with some modifications. Briefly, bacterial strain was incubated with AIO6 (5.66 - 7250U/mL). After staining the adhered cells were stained with 0.1% crystal violet solution and solubilized with ethanol. Then OD_595_ was measured using a microtiter plate reader (Synergy H1, BioTek, USA).

For structural analysis of the biofilm after treatment with AIO6, the biofilm was grown using the methodology described above. The biofilm was stained by adding Acridine orange (AO) / ethidium bromide (EB) stain (100 µg/ml, mixed in 1:1 ratio). The samples were fixed with 4% paraformaldehyde and imaged with a fluorescence microscope (Leica DMIL LED, Leica Microsystems, Wetzlar, Germany).

#### RNA Extraction and Gene Expression Measurement

RNA was extracted with RNAprep pure Cell/Bacteria Kit and reverse-transcribed into cDNA using FastKing gDNA Dispelling RT SuperMix. Real-time PCR (*q*PCR) was realized using LightCycler 480 (Roche Diagnostics, Mannheim, Germany), and a FastFire *q*PCR PreMix (SYBR Green) kit. The primers used are indicated in Table S4. The resulting data for each gene were normalized using housekeeping gene *16s rDNA* and analyzed using the 2^−ΔCt^ method.

#### Gut microbiota analysis

At the end of the feeding trial, the gut contents of 2-month-old zebrafish were collected 4 h post feeding. The gut contents were collected under aseptic conditions. Each replicate specimen was pooled gut content sample from six fish. Bacteria DNA was extracted and the bacterial V3-V4 region was amplified using barcode-indexed primers (338F: 5′-ACTCCTACGGGAGGCAGCA-3′ and 806R: 5′-GGACTACHVGGGTWTCTAAT-3′). PCR products were sequenced using the Illumina MiSeq platform and analyzed on the Majorbio Cloud platform (https://cloud.majorbio.com). Microbiota sequencing data have been deposited with the NCBI BioProject database (http://www.ncbi.nlm.nih.gov/bioproject) under Bioproject ID: PRJNA1209212.

The number of total bacteria or a specific phylum/genus was quantified by *q*PCR. Primers have been described^66–68^ and listed in Table S4. For 2-month-old zebrafish, results were expressed as Log_10_ copy numbers of bacterial 16S rDNA per milligram of gut contents. The samples of gnotobiotic zebrafish were collected under sterile conditions and washed three times in sterile PBS. Then ∼20 fish were collected and extracted DNA for *q*PCR quantification, the resulting data for each species were normalized using the data obtained by the gene primers for universal bacteria. For mono-associated zebrafish, the remaining samples were also homogenized in sterile PBS under sterile conditions. The bacterial numbers were determined by plating tenfold serial dilutions of the homogenates on LB agar plates supplemented with Amp, followed by incubation at 30L for 16 h. The Colony-forming units per zebrafish were counted. For the gut microbiota cultured *in vitro*, results were expressed as Log_10_ copy numbers of bacterial 16S rDNA per mL bacteria.

#### Gene knockdown

Gene knockdown was conducted using vivo morpholino oligonucleotides (MO) designed and synthesized by Gene Tools (Philomath, OR). The sequences of MO used in this study are as

follows: MyD88 MO, 5′-ATATCCACAAAGCAACATGCCTTTT-3′; spi1b MO, MO1 5′-CCTCCATTCTGTACGGATGCAGCAT-3′, MO2 5′-GGTCTTTCTCCTTACCATGCTCTCC-3′; Csf3r MO, 5′-AAGCACAAGCGAGACGGATGCCAT-3′; and control MO, 5′-CCTCTTACCTCAGTTACAATTTATA-3.′

#### Preparation and treatment of cell free supernatant and cell lysate of *P. shigelloides*

*A. P. shigelloides* CB5 was cultured in LB medium at 30L for 16 h. Cell free supernatant was obtained by centrifugation at 10, 000 g for 5 min and filtration through a 0.22-µm filter (Millipore, Darmstadt, Germany). The pellet was washed three times with PBS, and resuspended in same volume of PBS. The cell lysate was obtained by ultrasonication, centrifugation and filtration. For heat treatment experiments, cell free supernatant of *P. shigelloides* was boiled at 100L for 5 min and filtered through a 0.22-µm filter. For experiments using proteinase K, cell free supernatant of *P. shigelloides* was incubated with protease K at 70°C for 10 min. When applicable, CB5 was cultured in LB medium containing 10 μM GW4869 or an equal volume of vehicle (DMSO). CB5 was shaken at 30L and 180 rpm for 24 h, and the cell free supernatant was collected in the same way as described above. All samples were added to gnotobiotic zebrafish mono-associated with *Aeromonas* at 4 dpf by immersion.

#### *In vitro* interactions among *A. veronii* and *P. shigelloides*

A co-culture assay between *A. veronii* and *P. shigelloides* was performed as described previously.^69^ Briefly, at the start of the co-culture experiment, bacterial colony was cultured in LB medium overnight. Cells were centrifuged at 6, 000 g for 5 min, washed three times with PBS, and adjusted to an OD_600_ of 0.2. Then 50 μL of *A. veronii*, 50 μL of *P. shigelloides*, and 900 μL of LB medium were added to 24-well plates. The co-cultures were incubated at 30°C for 24 h. To measure changes in bacterial abundance, the abundance of *A. veronii* and *P. shigelloides* were performed using both *q*PCR and the plate count method. Compared with the growth of mono-culture, the fold changes in the growth of co-culture for each species were calculated. Meanwhile, we evaluated the potential of *A. veronii* and *P. shigelloides* to utilize various substrates using EcoPlate. *A. veronii* and *P. shigelloides* were cultured overnight and were evaluated by OD_600._ According to the manufacturer’s instructions, the pellet was washed three times with PBS, and resuspended in PBS. Then *A. veronii* and *P. shigelloides* were inoculated in 96-microplates and cultured for 72 h at 30L.

In order to evaluate whether AIO6 and the cell free supernatant of *P. shigelloides* directly inhibit the growth of *A. veronii*, *in vitro* culture experiment of *A. veronii* was conducted. Freshly grown *A. veronii* (1 × 10^7^ bacteria) were incubated alone or with the cell free supernatant of *P. shigelloides* alone or preincubated with 100 U/mL AIO6. After 24 h of the static culture at 30°C, bacterial cells were calculated by plating method.

#### Construction of *A. veronii* mutant

*A. veronii* XMX-5 Δ*ahyI*, Δ*ahyR,* Δ*lafK*, Δ*rpoN*, Δ*ascV*, Δ*exeMN*, Δ*minD*, Δ*vasH*, Δ*fliJ*, and Δ*flgB* mutant were constructed by double crossover allelic exchange using suicide vector pRE112.^70^ In brief, suicide vector pRE112 was digested with restriction enzymes SmaI, and upstream and downstream flanking regions corresponding to the target gene from *A. veronii* XMX-5 were PCR amplified (Table S5). The products were ligated into pRE112 to construct the knockout plasmids pRE112ΔahyI, pRE112ΔahyR, pRE112ΔlafK, pRE112ΔrpoN, pRE112ΔascV, pRE112ΔexeMN, pRE112ΔminD, pRE112ΔvasH, pRE112ΔfliJ, and pRE112ΔflgB. The knockout plasmid was then transformed into *E. coli* S17-1 (λpir) and then conjugated into *A. veronii*. The first homologous recombination integration was screened by growth on LB solid plates containing Cm and Amp. The double-crossover (DC) was screened on the plates containing Amp and 15% sucrose. The gene deletion mutant was confirmed by PCR and sequencing.

#### *In vitro* microbicidal assays of neutrophil elastase

The bactericidal activity of neutrophil elastase *in vitro* was evaluated. Briefly, *A. veronii* XMX-5 and *A. veronii* XMX-5Δ*flgB* were cultured at 30°C overnight and washed twice with PBS to remove any excess medium. Next, *A. veronii* (1 × 10^8^ bacteria cells) were incubated in the presence or absence of NE (2 μg) in a total volume of 200 μL PBS at 30°C. Following incubation for 2 h, viable counts were determined by plating method. After incubation for 8 h, the bacterial morphology was observed by scanning electron microscopy (SEM, SU-8010, HITACHI, Japan).

#### Statistical analysis

All data were performed using GraphPad Prism 9 software (GraphPad Software Inc. CA, USA). All data were expressed as mean ± SD. Comparisons between the two groups were analyzed using the unpaired Student’s *t*-test. Statistical significance was denoted in figures as \**p* < 0.05, \*\**p* < 0.01, \*\*\**p* < 0.001.

