## Supplementary Information for "Dietary quorum quenching AHL lactonase impairs the adaptation of commensal bacterium to host innate immunity"

**Supplementary Figures**

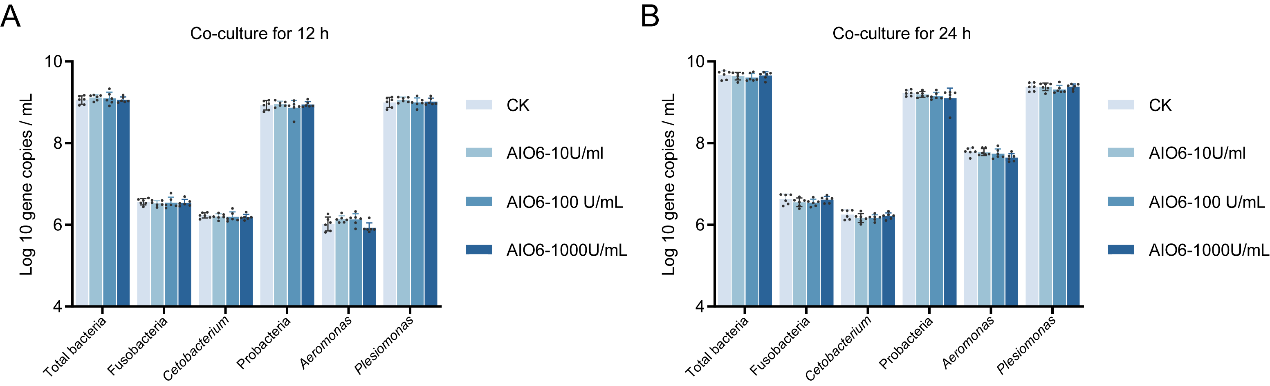

**Figure S1. AIO6 does not affect the microbiota composition under *in vitro* condition.**

(A and B) The microbiota from the gut of adult zebrafish was cultured for 12 h (A) and 24 h (B) in M9 (minimal medium) supplemented with PBS, 10 U/mL, 100 U/mL, or 1000 U/mL AIO6, respectively (n = 6).

Data are expressed as the mean ± SD.

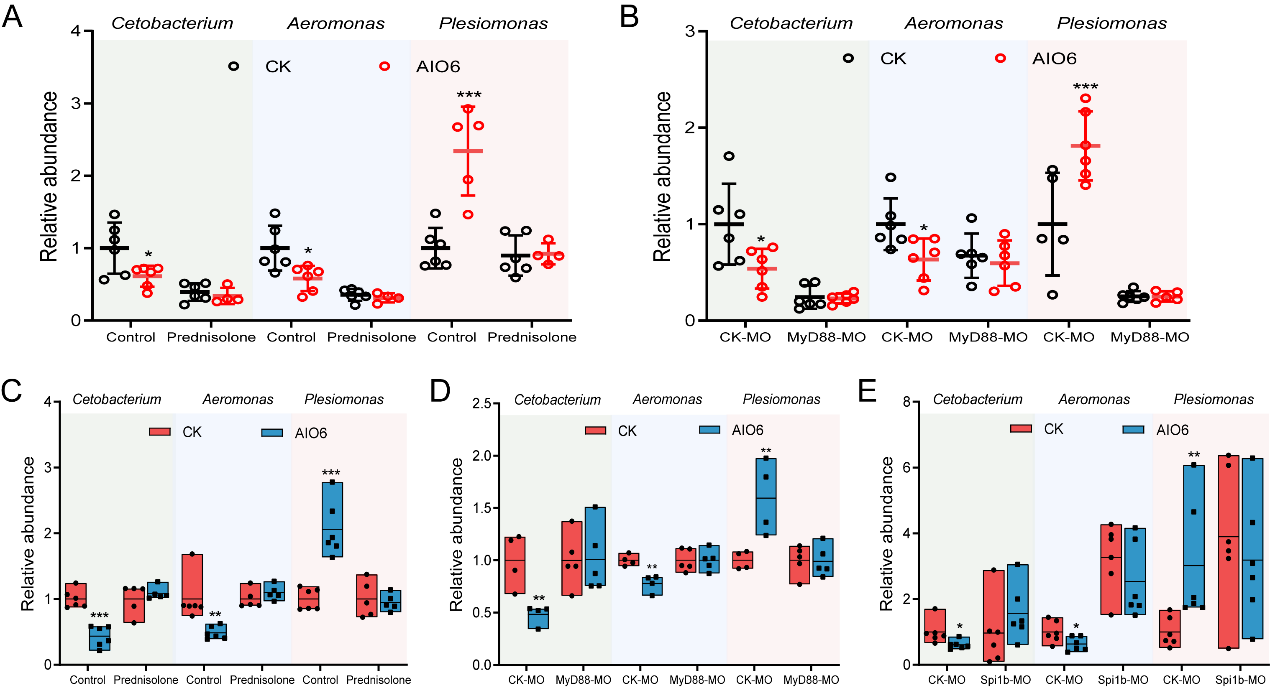

**Figure S2. Immunity inhibition blocked the effect of AIO6 on the commensal bacteria in conventional zebrafish and gnotobiotic zebrafish colonized with triple species.**

(A and B) Effect of prednisolone, or MyD88 MO on the relative abundance of *Aeromonas*, *Plesiomonas*, and *Cetobacterium* in conventional zebrafish larvae after AIO6 feeding (n = 6, pool of 20 zebrafish per sample).

(C-E) Effect of prednisolone, MyD88 MO, or Spi1b MO on the relative abundance of *Aeromonas*, *Plesiomonas*, and *Cetobacterium* in gnotobiotic zebrafish after AIO6 feeding (n = 5 or 6, pool of 20 zebrafish per sample).

Data are expressed as the mean ± SD. **p* < 0.05, ***p* < 0.01, ****p* < 0.001.

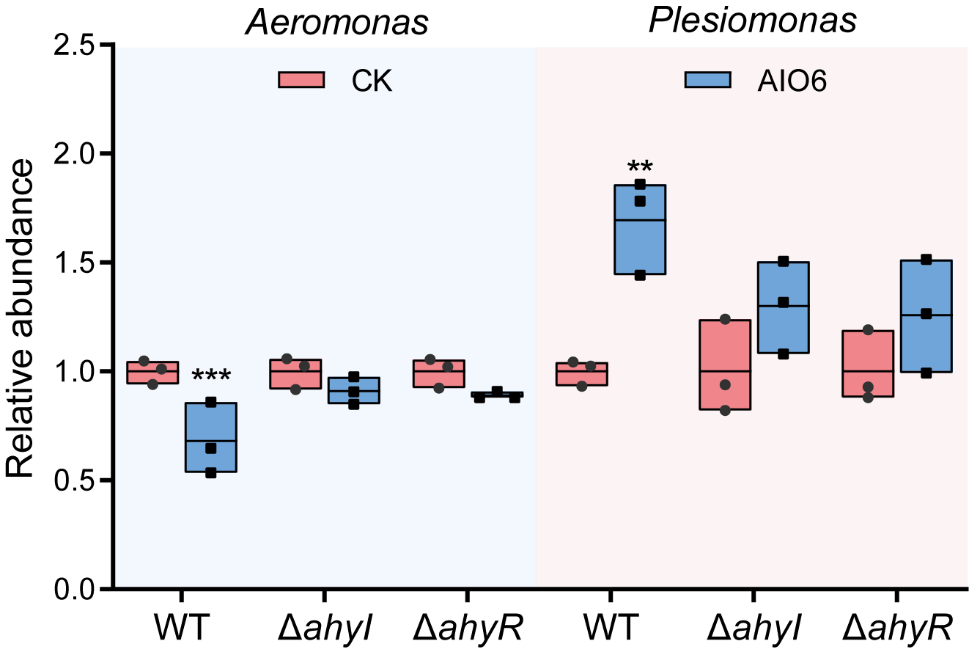

**Figure S3.** The relative bacterial abundance in gnotobiotic zebrafish colonized with *Plesiomonas* and *Aeromonas* WT strain or mutant deficient in the genes related to AHL synthesis/reception (n = 3, pool of 20 zebrafish per sample)*.* **p* < 0.05, ***p* < 0.01.

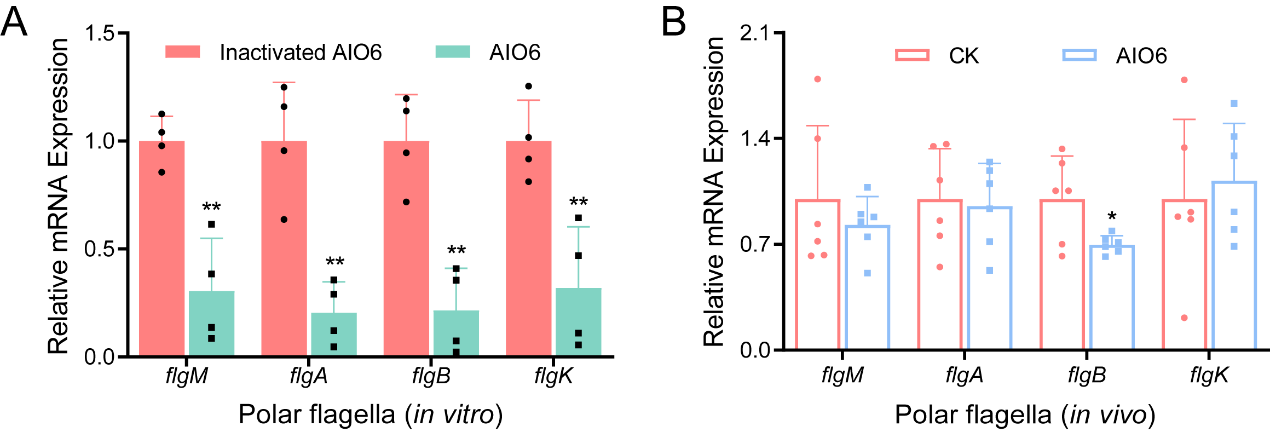

**Figure S4. AIO6 inhibited the expression of genes related to polar flagella of commensal *Aeromonas* under *in vitro* and *in vivo* conditions.**

(A and B) Gene expression of the polar flagellar system of commensal *Aeromonas* strain co-cultured with AIO6 under *in vitro* condition (A, n = 4) or in mono-associated gnotobiotic zebrafish fed AIO6 (B, n = 6, pool of 25 zebrafish per sample).

Data are expressed as the mean ± SD. **p* < 0.05, ***p* < 0.01, ****p* < 0.001.

**Supplementary Table 1**. Feed formulation and chemical composition of diets for adult zebrafish (g/kg, air dry basis)

| Ingredients | Groups | | |
| --- | --- | --- | --- |
|  | Control diet | AIO6 diet | AidB diet |
| Casein | 400.00 | 400.00 | 400.00 |
| Gelatin | 100.00 | 100.00 | 100.00 |
| Dextrin | 280.00 | 280.00 | 280.00 |
| Soybean oil | 60.00 | 60.00 | 60.00 |
| Lysine | 3.30 | 3.30 | 3.30 |
| VC phosphate | 1.00 | 1.00 | 1.00 |
| Vitamin premix^1^ | 2.00 | 2.00 | 2.00 |
| Mineral premix^2^ | 2.00 | 2.00 | 2.00 |
| Monocalcium phosphate | 20.00 | 20.00 | 20.00 |
| Choline chloride | 2.00 | 2.00 | 2.00 |
| Sodium alginate | 20.00 | 20.00 | 20.00 |
| Microcrystalline cellulose | 40.00 | 40.00 | 40.00 |
| Zeolite powder | 69.70 | 69.68 | 69.68 |
| AIO6 | 0.00 | 0.02 | 0.00 |
| AidB | 0.00 | 0.00 | 0.02 |
| Total | 1000.00 | 1000.00 | 1000.00 |
| Proximate analysis | | | |
| Crude protein | 42.00 | 42.00 | 42.00 |
| Crude fat | 6.01 | 6.01 | 6.01 |
| Gross energy, KJ/g | 14.02 | 14.02 | 14.02 |

^1^Containing the following (g/kg vitamin premix): thiamine, 0.438; riboflavin, 0.632; pyridoxine⋅HCl, 0.908; *D*-pantothenic acid, 1.724; nicotinic acid, 4.583; biotin, 0.211; folic acid, 0.549; vitamin B12, 0.001; inositol, 21.053; menadione sodium bisulfite, 0.889; retinyl acetate, 0.677; cholecalciferol, 0.116; *DL*-α-tocopherol-acetate, 12.632.

^2^Containing the following (g/kg mineral premix): CoCl_2_⋅6H_2_O, 0.074; CuSO_4_⋅5H_2_O, 2.5; FeSO_4_⋅7H_2_O, 73.2; NaCl, 40.0; MgSO_4_⋅7H_2_O, 284.0; MnSO_4_⋅H_2_O, 6.50; KI, 0.68; Na_2_SeO_3_, 0.10; ZnSO_4_⋅7H_2_O, 131.93; Cellulose, 501.09.

**Supplementary Table 2**. Feed formulation and chemical composition of diets for adult zebrafish (g/kg, air dry basis)

| Ingredients | Groups | |
| --- | --- | --- |
|  | Control diet | Antibiotic diet |
| Casein | 400.00 | 400.00 |
| Gelatin | 100.00 | 100.00 |
| Dextrin | 280.00 | 280.00 |
| Soybean oil | 60.00 | 60.00 |
| Lysine | 3.30 | 3.30 |
| VC phosphate | 1.00 | 1.00 |
| Vitamin premix^1^ | 2.00 | 2.00 |
| Mineral premix^2^ | 2.00 | 2.00 |
| Monocalcium phosphate | 20.00 | 20.00 |
| Choline chloride | 2.00 | 2.00 |
| Sodium alginate | 20.00 | 20.00 |
| Microcrystalline cellulose | 40.00 | 40.00 |
| Zeolite powder | 69.70 | 63.90 |
| Polymyxin B | 0.00 | 2.50 |
| Neomycin | 0.00 | 3.30 |
| Total | 1000.00 | 1000.00 |
| Proximate analysis | | |
| Crude protein | 42.00 | 42.00 |
| Crude fat | 6.01 | 6.01 |
| Gross energy, KJ/g | 14.02 | 14.02 |

^1^Containing the following (g/kg vitamin premix): thiamine, 0.438; riboflavin, 0.632; pyridoxine⋅HCl, 0.908; *D*-pantothenic acid, 1.724; nicotinic acid, 4.583; biotin, 0.211; folic acid, 0.549; vitamin B12, 0.001; inositol, 21.053; menadione sodium bisulfite, 0.889; retinyl acetate, 0.677; cholecalciferol, 0.116; *DL*-α-tocopherol-acetate, 12.632.

^2^Containing the following (g/kg mineral premix): CoCl_2_⋅6H_2_O, 0.074; CuSO_4_⋅5H_2_O, 2.5; FeSO_4_⋅7H_2_O, 73.2; NaCl, 40.0; MgSO_4_⋅7H_2_O, 284.0; MnSO_4_⋅H_2_O, 6.50; KI, 0.68; Na_2_SeO_3_, 0.10; ZnSO_4_⋅7H_2_O, 131.93; Cellulose, 501.09.

**Supplementary Table 3**. Feed formulation and chemical composition of diets for larval zebrafish (g/kg, air dry basis)

| Ingredients | Groups | |
| --- | --- | --- |
|  | Control diet | AIO6 diet |
| Casein | 460.00 | 460.00 |
| Gelatin | 110.00 | 110.00 |
| Dextrin | 180.00 | 180.00 |
| Lard oil | 30.00 | 30.00 |
| Soybean oil | 30.00 | 30.00 |
| Fish liver oil | 20.00 | 20.00 |
| Soybean lecithin | 20.00 | 20.00 |
| Lysine | 1.80 | 1.80 |
| VC phosphate | 1.00 | 1.00 |
| Vitamin premix^1^ | 2.00 | 2.00 |
| Mineral premix^2^ | 2.00 | 2.00 |
| Monocalcium phosphate | 20.00 | 20.00 |
| Choline chloride | 2.00 | 2.00 |
| Sodium alginate | 20.00 | 20.00 |
| Zeolite | 101.20 | 101.18 |
| AIO6 | 0.00 | 0.02 |
| Total | 1000 | 1000 |
| Proximate analysis | | |
| Crude protein | 48.09 | 48.09 |
| Crude fat | 9.90 | 9.90 |
| Gross energy, KJ/g | 18.60 | 18.60 |

^1^Containing the following (g/kg vitamin premix): thiamine, 0.438; riboflavin, 0.632; pyridoxine⋅HCl, 0.908; *D*-pantothenic acid, 1.724; nicotinic acid, 4.583; biotin, 0.211; folic acid, 0.549; vitamin B12, 0.001; inositol, 21.053; menadione sodium bisulfite, 0.889; retinyl acetate, 0.677; cholecalciferol, 0.116; *DL*-α-tocopherol-acetate, 12.632.

^2^Containing the following (g/kg mineral premix): CoCl_2_⋅6H_2_O, 0.074; CuSO_4_⋅5H_2_O, 2.5; FeSO_4_⋅7H_2_O, 73.2; NaCl, 40.0; MgSO_4_⋅7H_2_O, 284.0; MnSO_4_⋅H_2_O, 6.50; KI, 0.68; Na_2_SeO_3_, 0.10; ZnSO_4_⋅7H_2_O, 131.93; Cellulose, 501.09.

**Supplementary Table 4.** Sequences of primers used for *q*PCR analysis

| Primer | Forward (5'→3') | Reverse (5'→3') |
| --- | --- | --- |
| Universal bacteria | CCTACGGGAGGCAGCAG | ATTACCGCGGCTGCTGG |
| Fusobacterium  (phylum) | KGGGCTCAACMCMGTATTGCGT | TCGCGTTAGCTTGGGCGCTG |
| Proteobacteria  (phylum) | TCGTCAGCTCGTGTYGTGA | CGTAAGGGCCATGATG |
| *Cetobacterium* (genus) | AGTTTGATCCTGGCTCAGGATG | GAGGCAAGTTCCTTACGCGTT |
| *Plesiomonas* (genus) | CTCCGAATACCGTAGAGTGCTATCC | CTCCCCTAGCCCAATAACACCTAAA |
| *Aeromonas* (genus) | GCTGTGTCCTTGAGACGTGGC | TTCTGATTCCCGAAGGCACTCC |
| 16s rDNA | CCTACGGGAGGCAGCAG | ATTACCGCGGCTGCTGG |
| aerolysin | GTCTGTGCCAGTGGTTATCGT | GCGAGATAGCTGACTGGTTTG |
| L-fliM | CGAGAAGTCCGCTTGAACCACAG | GTTCGGTATCGGTCAGCCCTTTG |
| L-fliA | AGCAAGCGAGCCGTCAATGTG | CGATGGCGACCGTATCAACTGG |
| L-fliE | CAGTCAGACACCGTCCATCGAATC | TCGTCACTCTGTCCCGCATCG |
| L-flhA | GGGAGTGATTGCTCGGGATAAACC | GGGCGGGTAGAGGTCTCGTTAC |
| L-flgB | GGATGGTAATACCGCTGAGTTGGG | TGAGGAAGGTGAGGCTGGTCTG |
| L-flgE | AGCCGAGTCAGCCAGTATGCC | ACCGCCGTCTCCATCCACAC |
| P-flgM | GGCAGCAAGTCAGTCACCTCATC | TTGTTGCGCCTCACTGGTCAAG |
| P-flgA | TCCGTTTGAGCCAGGTTTGTGTC | CTCTCCGAAGCTGCCATCTTGC |
| P-flgB | GCTACGCTCCTGCTGCACATC | AGTCGCATCAGGGGTTTGGTTTG |
| P-flgK | ATTGAAATGCCGTGACCCGTGAG | CTCGCTGGACAAGTCTGGTAATCG |

**Supplementary Table 5.** Primers used for the mutagenesis of *A. veronii* XMX-5 genes

| Primer | Sequence (5'→3') | Product Size (bp) |
| --- | --- | --- |
| ahyI-up-F | TAAGTGAACTGCATGaattcccGCGATGGTAGCTCCTTCTTAGAAATTCA | 1088 |
| ahyI-up-R | GTTTGATAGTGTCCGATAAGTTTCTCTACGGCGTTTCGTGGGTGA |  |
| ahyI-down-F | CACCCACGAAACGCCGTAGAGAAACTTATCGGACACTATCAAACTGCCAC | 1095 |
| ahyI-down-R | catgcgatatcgagctctcccGACTGAGGTTGAGTAGTGACCCGCCAG |  |
| ahyR-up-F | TAAGTGAACTGCATGaattcccGGCCGCGGGCCAACCAGGGC | 824 |
| ahyR-up-R | AACATTTCACTTCGGTGACGGATGCCGAATCTTGAACAGGTTGTTGTCACC |  |
| ahyR-down-F | ACAACCTGTTCAAGATTCGGCATCCGTCACCGAAGTGAAATGTTCCAGAT | 860 |
| ahyR-down-R | catgcgatatcgagctctcccAGCCCTGCGGGGTCATAGCGATACAGC |  |
| lafK-up-F | TAAGTGAACTGCATGaattcccgctgttggataaatgcgctgatggggaa | 706 |
| lafK-up-R | aatgccttgctcacgcatcgccgcgagcccttcaaactcatgcagaatacgtg |  |
| lafK-down-F | tgcatgagtttgaagggctcgcggcgatgcgtgagcaaggcattgatatt | 641 |
| lafK-down-R | catgcgatatcgagctctccctgccatacaggggcacatagtttttgctgg |  |
| rpoN-up-F | TAAGTGAACTGCATGaattcccatcaggacatcagcctgcagcccat | 705 |
| rpoN-up-R | aaggtgggataaacaaggattcgcgtaattgcggcgtcatggtcagggactga |  |
| rpoN-down-F | tccctgaccatgacgccgcaattacgcgaatccttgtttatcccaccttccaatc | 654 |
| rpoN-down-R | gcatgcgatatcgagctctccccaaaggtcatcaggatcgcggttggcgg |  |
| exeMN-up-F | TAAGTGAACTGCATGaattccctcagccctgcaactcggcaatgagtg | 837 |
| exeMN-up-F | ccttgctgcatctgtggtggcaaccaaccgctgttcccgtgctgtgat |  |
| exeMN-down-F | agcacgggaacagcggttggttgccaccacagatgcagcaagggct | 828 |
| exeMN-down-R | catgcgatatcgagctctccccatctttgaaaccagcagtgaagtggacatc |  |
| ascV-up-F | TAAGTGAACTGCATGaattcccatctccctagacgatcaggagcgaagc | 741 |
| ascV-up-R | tgggtcagctcctgataggagatgtccttgcgctcgccgatccg |  |
| ascV-down-F | gatcggcgagcgcaaggacatctcctatcaggagctgacccagcagatca | 731 |
| ascV-down-R | catgcgatatcgagctctcccagctgacaggtctgatccgagagggcta |  |
| vasH-up-F | TAAGTGAACTGCATGaattcccGccatctatcaaatctccgactcggtgca | 1218 |
| vasH-up-R | aagatgctggcgagtgtgctgcagggattgccaccctctatcgcaagataaag |  |
| vasH-down-F | ttgcgatagagggtggcaatccctgcagcacactcgccagcatcttggcaa | 687 |
| vasH-down-R | catgcgatatcgagctctcccacatagccatcggcggcatcggtattct |  |
| minD-up-F | TAAGTGAACTGCATGaattcccccggattggtgcccatgaacagctgc | 632 |
| minD-up-R | ttccaggaaaaactccagatggacgcattttgcgcagaccactcgcctgat |  |
| minD-down-F | ggcgagtggtctgcgcaaaatgcgtccatctggagtttttcctggaaaatttgttgc | 641 |
| minD-down-R | catgcgatatcgagctctcccatctgaaatcgctcggcagccaattc |  |
| fliJ-up-F | TAAGTGAACTGCATGaattcccagtcggcaagagtatgttgctgggcat | 932 |
| fliJ-up-R | aatcttggcgctgagcgtcaaggcgagatcaactgcagctggatga |  |
| fliJ-down-F | ccagctgcagttgatctcgccttgacgctcagcgccaagattttccag | 971 |
| fliJ-down-R | gcatgcgatatcgagctctccccttagctcaggcaagcagatacaggcat |  |
| flgB-up-F | TAAGTGAACTGCATGaattcccatgcctgtatctgcttgcctgagctaag | 915 |
| flgB-up-R | gaggaaggtgaggctggtctggccctgacatcgagggcataagggtgta |  |
| flgB-down-F | ttatgccctcgatgtcagggccagaccagcctcaccttcctcaatatgaa | 952 |
| flgB-down-R | catgcgatatcgagctctcccgtaccttgcaactcgatctgaccgc |  |
